# SCOPE: Flexible targeting and stringent CARF activation enables type III CRISPR-Cas diagnostics

**DOI:** 10.1101/2021.02.01.429135

**Authors:** Jurre A. Steens, Yifan Zhu, David W. Taylor, Jack P.K. Bravo, Stijn H.P Prinsen, Cor D. Schoen, Bart J.F Keijser, Michel Ossendrijver, L. Marije Hofstra, Stan J.J. Brouns, Akeo Shinkai, John van der Oost, Raymond H.J. Staals

**Affiliations:** Laboratory of Microbiology, Wageningen University and Research, 6708 WE, Wageningen, The Netherlands; Department of Molecular Biosciences, University of Texas at Austin, Austin, TX, USA; Institute for Cellular and Molecular Biology, University of Texas at Austin, Austin, TX 78712, USA; Center for Systems and Synthetic Biology, University of Texas at Austin, Austin, TX 78712, USA; LIVESTRONG Cancer Institutes, Dell Medical School, Austin, TX 78712, USA; Scope Biosciences, Bronland 10, 6708 WH Wageningen, The Netherlands; BioInteractions and Plant Health, Wageningen Plant Research, P.O. Box 69, 6700 AB, Wageningen, The Netherlands; TNO, Utrechtseweg 48, 3704 HE, Zeist, The Netherlands; Virology, Department of Medical Microbiology, University Medical Center Utrecht, 3584 CX Utrecht, The Netherlands; Department of Bionanoscience, Delft University of Technology, van der Maasweg 9, 2629 HZ Delft, The Netherlands; RIKEN SPring-8 Center, Sayo, Hyogo 679-5148, Japan; RIKEN Cluster for Pioneering Research, Wako, Saitama 351-0198, Japan

**Author notes:** These authors contributed equally (Jurre A. Steens, Yifan Zhu).

**Keywords:** CRISPR-Cas, type III, Cas10, seed, cyclic oligoadenylates, nucleic acids detection, SARS-CoV-2, point-of-care, Cmr complex, SCOPE

## Abstract

Characteristic properties of type III CRISPR-Cas systems include recognition of target RNA (rather than DNA) and the subsequent induction of a multifaceted immune response. This involves sequence-specific cleavage of a target RNA and production of cyclic oligoadenylate (cOA) second messenger molecules that may trigger dormancy or cell death. In this study, we discovered that a largely exposed seed region at the 3’ end of the crRNA is essential for target RNA binding and cleavage, whereas base pairing at a unique region at the 5’ end of the guide is required to trigger cOA production. Moreover, we uncovered that the natural variation in the composition of type III complexes within a single host results in different guide lengths, and hence variable seed regions. This shifting seed may prevent escape by invading genetic elements, while controlling cOA production very tightly to prevent unnecessary damage to the host. Lastly, we used these findings to develop a new diagnostic tool, named SCOPE, which was used for the specific detection of SARS-CoV-2 from human nasal swab samples, showing sensitivities in the atto-molar range.

## Introduction

As a widespread prokaryotic adaptive immune system, CRISPR-Cas (Clustered Regularly Interspaced Short Palindromic Repeats/CRISPR-associated) systems target and cleave genetic material of viruses and other mobile genetic elements (MGEs) [1]–[4]. A CRISPR array is composed of alternating repeat and spacer sequences, typically with the repeats consisting of identical sequences and the spacers consisting of variable sequence fragments acquired from invading MGEs [5]–[7]. The CRISPR array is generally located adjacent to a set of CRISPR-associated (cas) genes encoding the Cas proteins.

In type III CRISPR-Cas systems, the CRISPR array is expressed and processed into mature CRISPR RNA (crRNA) by the Cas6 ribonuclease [8]. A second maturation event occurs where the crRNA is trimmed at the 3’ end [9][10]. The matured and trimmed crRNAs form a ribonucleoprotein complex together with a set of Cas proteins: the type III effector complex (Figure 1A). In the interference stage, type III complexes specifically bind and degrade target RNA sequences (protospacers) complementary to the bound crRNA guides.

**Figure 1.**
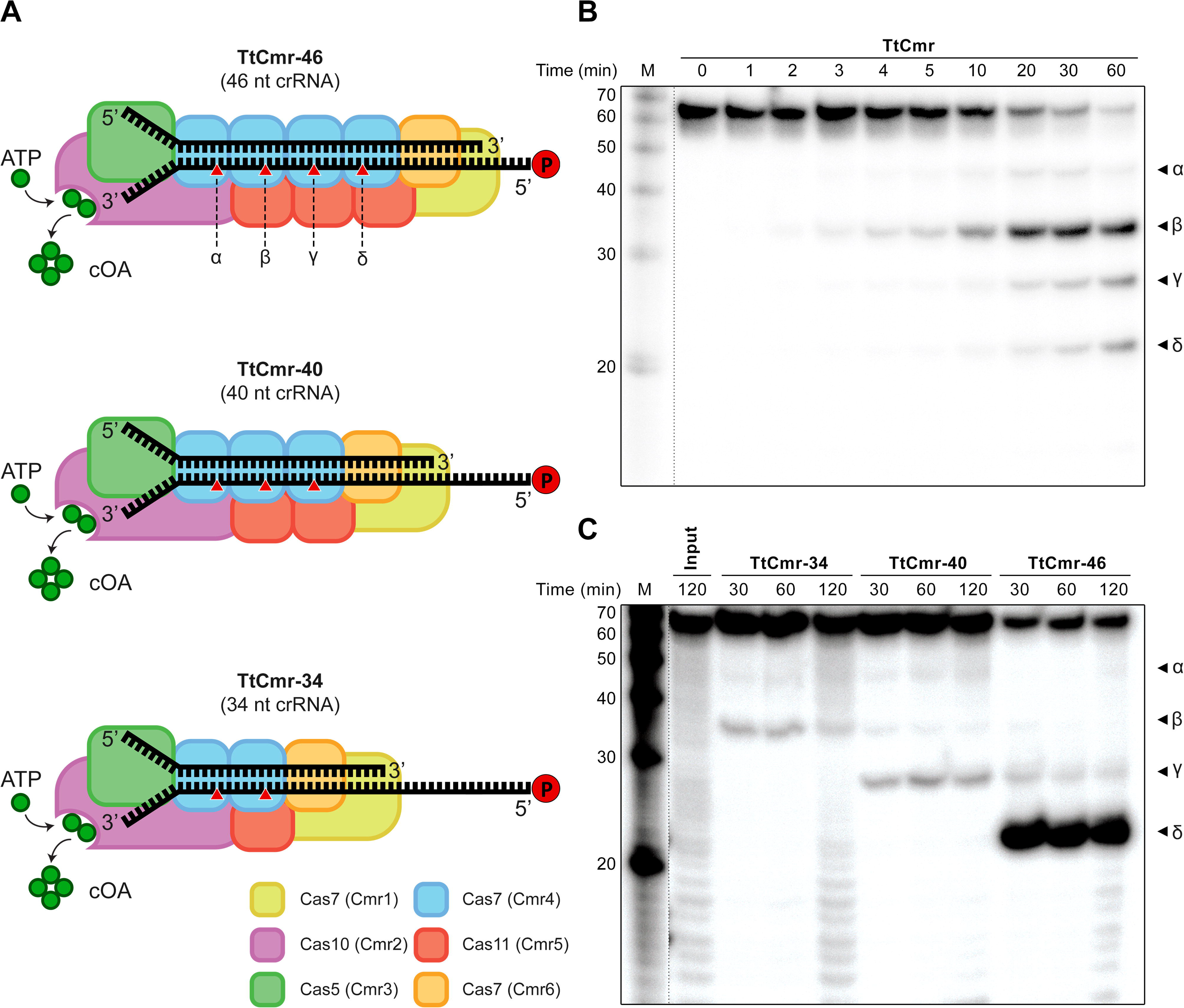
*In vitro* RNase activity assays with the endogenous and reconstituted TtCmr complexes. **(A)** Schematic illustration of the different reconstituted TtCmr complexes used in the activity assays shown in panel C, pre-loaded with either the 46 (TtCmr-46), 40 (TtCmr-40) or 34 nt (TtCmr-34) crRNA (top strand). Red triangles indicate the anticipated cleavage sites (α, β, γ and δ) in the 4.5 target RNA (bottom strand, table S1) by the endoribonuclease activity of the Cas7 subunits. The target RNA was radiolabeled at the 5’ end with ^32^P γ-ATP (“P” in the red circle) **(B)** Denaturing PAGE analysis of the activity assay using a 5’ labeled target RNA complementary to the crRNA incubated with the endogenous TtCmr complex. A single stranded RNA marker (“M”) was used as size standards as indicated on the left. **(C)** Activity assays similar to panel B but using the reconstituted complexes. Discontinuous gel lanes are indicated by a dashed line.

Unlike other CRISPR-Cas systems that exclusively target either DNA (type I, II and V) or RNA (type VI), many type III systems have the capacity to target both nucleic acids [11]–[15][16][17]. Structural and biochemical analyses of the multi-subunit type III complexes have revealed the subunits responsible for these activities and showed that they are induced in a step-wise manner, starting with the binding of a complementary target RNA [15][18]–[23]. Target RNA binding activates the RNase activity of Cas7, which is present in multiple copies and constitutes the backbone of type III complexes. As such, type III interference complexes have multiple (2-4) active sites, cleaving the target RNA at 6 nt intervals (Figure 1A) [12][15][18][24][25]. Simultaneously, target RNA binding activates two distinct catalytic domains of the Cas10 protein, the large subunit of type III complexes. Activation of the HD domain of Cas10 confers sequence-nonspecific DNase activity [14][16][26], while activation of its Palm domain triggers oligoadenylate cyclase activity, producing cyclic oligoadenylate (cOA) second messenger molecules that allosterically activate effector CARF (CRISPR-associated Rossmann fold) proteins [27][28][27][28]. Most of the CARF proteins that have been characterized so far appear to be promiscuous RNases, cleaving both viral and host RNAs, thereby potentially inducing cell dormancy or cell death [29][30][31][32].

Previously, we characterized the structural and enzymatic features of the endogenous type III-B Cmr complex from *T. thermophilus* HB8 (TtCmr) [23][18]. We showed that TtCmr adopts a structure similar to type I (Cascade) complexes: a Cas7 (Cmr4) backbone with multiple copies of the small subunit Cas11 (Cmr5). The complex is capped at one end by the large subunit Cas10 (Cmr2) and Cas5 (Cmr3) and is capped at the other end with two more Cas7-like subunits (Cmr1 and Cmr6) [23]. It is important to note that the Cas10 subunit of the TtCmr complex lacks the HD domain, and hence does not have DNase activity [18]. The mature crRNA runs along the Cas7 backbone of the complex, with its 5’ repeat-derived end (the 5’ handle) anchored by Cas10/Cas5, and its 3’ end located at the Cmr1/Cmr6 end [20][24][33]. Interestingly, the 3’ end of the mature crRNA is variable in type III systems, due to an uncharacterized 3’ processing event following the endonucleolytic cleavages of Cas6 [36][37]. Although the details of this 3’ processing event are not known, it is hypothesized that the heterogenous nature of the TtCmr complex might be responsible for this, as is explained below.

Firstly, analysis of the crRNA-content of the endogenous TtCmr complex showed that it indeed co-purifies with mature crRNAs of different sizes (with variable 3’ ends), with a distinct 6-nt pattern: 34, 40 & 46 nt [18]. Secondly, our cryo-EM structures revealed that the native population of TtCmr complexes consisted of larger (with a stoichiometry of Cmr1_1_2_1_3_1_4_4_5_3_6_1_) and smaller complexes (i.e. Cmr1_1_2_1_3_1_4_3_5_2_6_1_ and Cmr1_1_2_1_3_1_4_2_5_1_6_1_), with the smaller complexes lacking one or two Cas7–Cas11 (Cmr4-Cmr5) backbone segment(s) (Figure 1A). Taken together, these data strongly suggest that the 3’ end of the mature crRNA is determined by the stoichiometry of the TtCmr complex. In this scenario, it is likely that the 3’ end is generated by a (non-Cas) host ribonuclease, such as PNPase, that shortens the unprotected, protruding 3’ end of the bound crRNA [36].

In this study, we set out to understand the biological significance of these differently sized Cmr complexes. This has revealed an unexpected flexible seed region at the 3’ end of the crRNA guides, that might be important for these systems to prevent phage escapees. Additionally, we identified a fixed region at the 5’ end of the crRNA that triggers the catalytic activities of Cas10, thereby ensuring tight control over CARF protein activation. These characteristics formed the basis for the development of a highly-sensitive novel type III diagnostics platform called SCOPE (Screening using CRISPR Oligoadenylate-Perceptive Effectors).

## Results

### TtCmr complexes consist of a heterogenous mixture of distinct compositions

We previously demonstrated that endogenous type III-B Cmr complexes purified from *T. thermophilus* HB8 (TtCmr) are loaded with mature crRNA guides of different lengths (34-40-46 nt), and that the crRNA-4.5 (CRISPR array 4, spacer 5) is most abundant (Figure 1A)[18]. This endogenous Cmr complex specifically cleaves complementary target RNAs (4.5 target RNA) at 6 nt intervals, corresponding to the Cas7 subunits in the backbone of the complex. Consequently, this results in 5’ labeled degradation products of 39 (α), 33 (β), 27 (γ) and 21 (δ) nucleotides (Figure 1B). However, the heterogeneous nature of the crRNA-content of the endogenous Cmr complexes [18][23], complicates the interpretation of these results. Therefore, to further reveal the mechanism of target RNA cleavage, we used *E.coli*-produced subunits to reconstitute three different Cmr complexes bound to a single crRNA (crRNA-4.5) of a defined length. We chose to include crRNA lengths of either 34 (TtCmr-34), 40 (TtCmr-40) or 46 nt (TtCmr-46), based on their abundance in their native host [18]. Opposed to all four (α-δ) 5’ - labeled degradation products observed with the endogenous complex, each of the reconstituted complexes produced defined degradation products decreasing in size with a longer crRNA (Figure 1C). This is consistent with the idea that the composition of the complex corresponds to the length of the crRNA, with larger complexes (e.g. TtCmr-46) harboring more cleavage sites, hence cleaving more closely to the labeled 5’ end of the target RNA. Smaller complexes, such as TtCmr-40 and TtCmr-34, lack one or two Cas7–Cas11 (Cmr4-Cmr5) backbone segment(s), respectively, and therefore cleave the target RNA at less and more distal locations (further away from the 5’ label), resulting in larger degradation products. These results show that the population of endogenous TtCmr complexes is a heterogenous mixture of bigger and smaller complexes, cleaving their cognate target RNAs at different positions.

### A flexible 3’ seed region regulates target cleavage and cyclic oligoadenylate production

To investigate the significance of these type III complexes with different stoichiometries, we performed activity assays to probe for differences in seed requirements for RNA targeting as well as for the production of cyclic oligoadenylate (cOA) second messengers. In the structurally similar type I effector complexes (i.e. the Cascade complex), DNA targeting is governed by two factors: the PAM (protospacer adjacent motif) and the seed [37]–[39]. In the RNA targeting type III systems, however, self/non-self discrimination is conferred by an rPAM (RNA protospacer-adjacent motif) [14]. This motif checks for complementarity between the 5’ handle of the crRNA (8 nucleotides, referred to as nucleotides −8 to −1) and the corresponding 3’ region flanking the protospacer [14][16][42] (Figure 2A). Since TtCmr is devoid of DNase activity [18][23] we tested whether RNase activity and production of cOA are affected by target RNAs with complementarity to the 5’ handle of the crRNA (Figure 2A). The cleavage activity assays with the endogenous Cmr complex showed that these ‘self-like’ substrates had no substantial effect on RNA cleavage activity (Figure 2B). To probe for their impact on cOA production, we developed a pyrophosphatase-based assay that directly reflects the oligoadenylate oligomerization, as a measure for cOA production. A by-product of cOA production is the formation of pyrophosphate (PPi), which can be converted to free phosphate (Pi) by a thermostable pyrophosphatase enzyme [27]. This Pi can be visualized by a range of colorimetric and fluorometric techniques, such as the Malachite Green colorimetric technique used in this study. Using this assay, we show that target RNAs with complementarity to the 5’ handle reduced the production of cOA to background levels, comparable to using a non-target (NT) target RNA (Figure 2C). Similar results were obtained for the reconstituted TtCmr-46 and TtCmr-40 complexes (Figure S1). We conclude that cOA production, but not target RNA cleavage, is affected by complementarity between target RNA and the 5’ handle of the crRNA, potentially due to mitigate detrimental consequences that cOA production might invoke (e.g. cell death/dormancy) when binding antisense transcripts from the CRISPR array [12][13].

**Figure 2.**
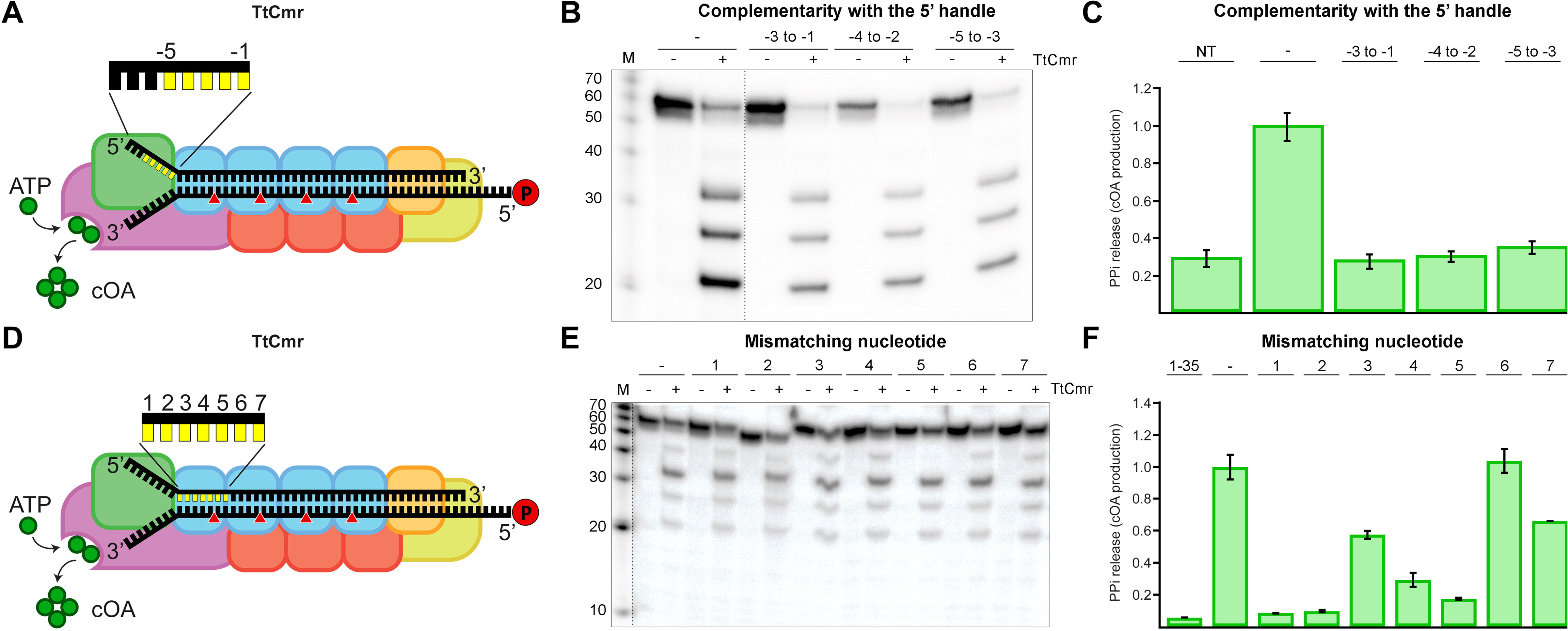
Impact of complementarity in the 5’ handle and mismatches in the first spacer region on RNA targeting and cOA production. **(A)** Schematic illustration of the TtCmr complex bound to a target RNA (4.5 target RNA, table S1), showing the different subunits in different colors, crRNA (top strand) and target RNA (bottom strand). The target RNA was labeled at the 5’ end with ^32^P (“P” in the red circle). Red triangles indicate the cleavage sites within TtCmr. Highlighted in yellow are the first 5 nucleotides in the 5’ handle of the crRNA (nucleotides −1 to −5). **(B)** Different target RNAs (table S1) with matches to the 5’ handle were used in an activity assay and analyzed on denaturing PAGE. **(C)** Impact of 5’ handle complementarity on PPI release i.e. COA production. **(D)** Similar schematic illustration as panel A with the first 7 nucleotides of the spacer region of the crRNA highlighted in yellow. **(E)** Similar activity assay as in panel B but with RNA targets (table S1) with single mismatches in the first 7 nucleotides of the spacer region of the crRNA. **(F)** Similar activity assay as panel E but depicting the release of PPI i.e. production of cOA. Discontinuous gel lanes are indicated by a dashed line.

Next, we tested whether TtCmr utilizes a seed similar the Cascade complexes of type I systems. Activity assays were performed by incubating the TtCmr complex with target RNAs containing single mismatch mutations in the first 7 nt of the spacer region of the crRNA (i.e. the region on the crRNA base pairing with the protospacer) (Figure 2D). The results showed that RNA targeting was not affected by these mutations, although a mismatch at position 5 abolished cleavage at the adjacent site, as demonstrated by the missing 39 nt degradation product (Figure 2E). Interestingly however, cOA production was greatly affected by these mismatches, in particular at positions 1 and 2 (Figure 2F). Similar results were obtained with the reconstituted TtCmr-46 and TtCmr-40 complexes (Figure S2). These results indicate that the seed is either lacking or located in a different region of the crRNA. However, since base pairing at the most 5’ region does appear to be critical for cOA production, we designated this segment as the Cas10-activating region (CAR).

Since the seed is defined as the region on the crRNA that initiates base pairing with its target, we performed EMSA binding assays with the endogenous TtCmr complex (Figure S3). To probe for regions crucial for initiating base paring, we used RNA targets with different mismatching segments (Figure 3A). We observed that targets with mismatches in the first three segments (nucleotides 1-5, 7-11, and 13-17) did not influence the binding of the target RNA by the TtCmr complex as the migration was similar to that of the fully complementary RNA target control (WT). However, mismatches in the fourth and fifth segments (nucleotides 19-23 and 25-29) substantially affected the electrophoretic mobility of the TtCmr/crRNA-target RNA tertiary complex, suggesting a seed region at the 3’ end of the crRNA.

**Figure 3.**
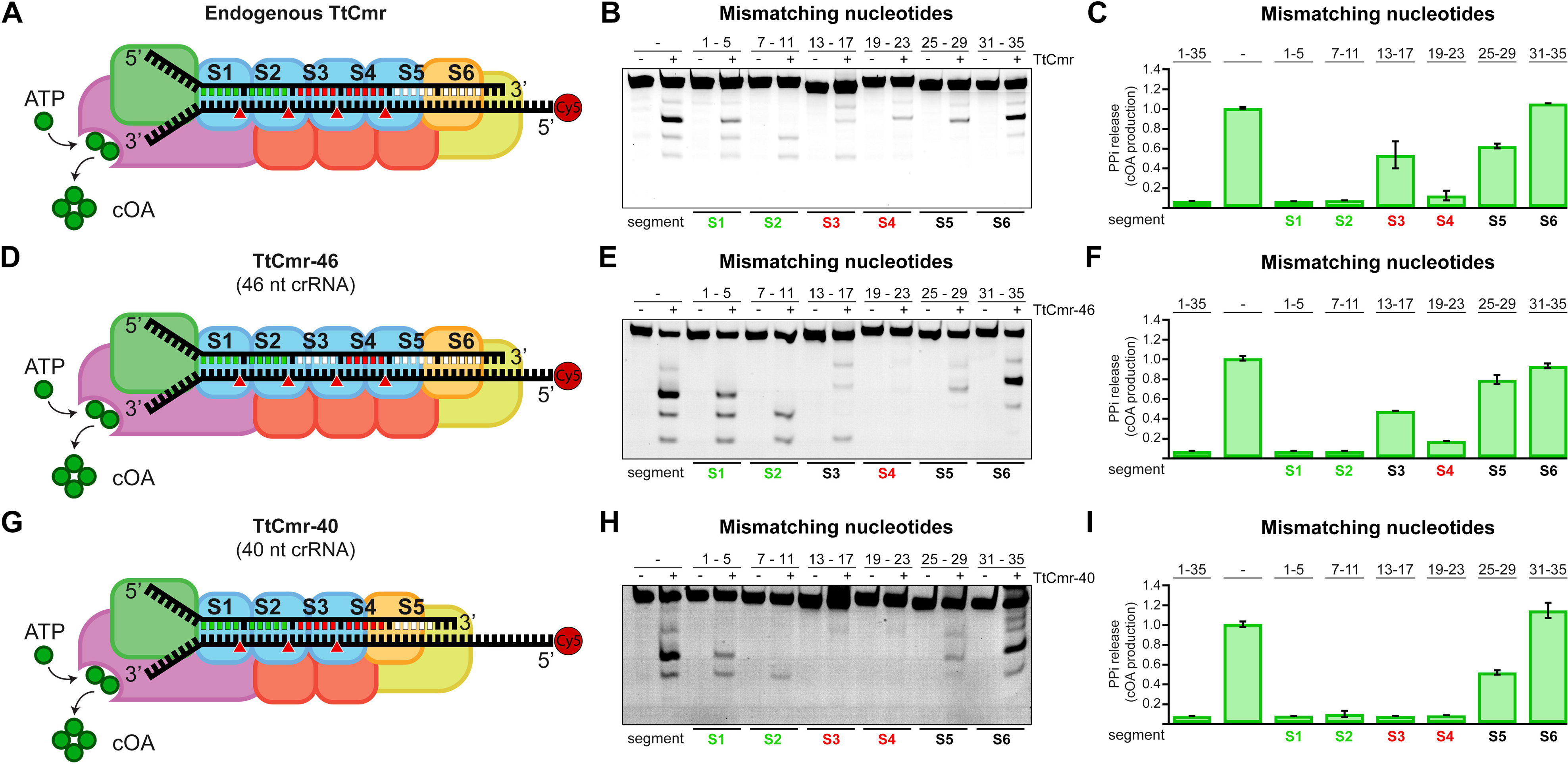
A flexible seed region at the 3’ end of the crRNA. **(A)** Schematic illustration of the endogenous TtCmr complex. **(B)** Different target RNAs with segments mismatches were used in an activity assay and analyzed on denaturing PAGE. **(C)** Impact of target RNAs with mismatches in the indicated segments on the production of cOA. **(D)** Schematic overview of the 46 nt crRNA complex (TtCmr-46). (E) Similar to panel B, using the 46 nt crRNA complex (TtCmr-46). **(F)** Similar to panel C, using 46 nt crRNA complex (TtCmr-46). **(G)** Schematic overview of the 40 nt crRNA complex (TtCmr-40). **(H)** Similar to panel B, using the 40 nt crRNA complex (TtCmr-40). **(I)** Similar to panel C, using the 40 nt crRNA complex (TtCmr-40). Target RNA contain a 5’ end Cy5 label (red circle); mismatched segments are indicated with S1-S6. Red triangles indicate the cleavage sites within TtCmr. CAR segments are indicated in green, seed segments are indicated in red.

To further investigate the impact of such a uniquely located seed on RNA degradation and cOA production, we performed activity assays with the endogenous TtCmr complexes using RNA targets containing different mismatching segments (Figure 3A-C). In agreement with our previous findings, RNA targets with mismatches in the first segment (S1, nucleotides 1-5) did not interfere with target degradation, despite skipping one cleavage site downstream of the mismatched segment. Similarly, mismatching of the segments S2-S5 (nucleotides 7-12, 13-17, 19-23 and 25-29) resulted in skipping both the adjacent (up- and downstream) cleavage sites, whereas cleavage at the other sites was unaffected. Mismatches in segment S6 (nucleotides 31-35) had no effect on RNA degradation, other than skipping the upstream cleavage site (Figure 3B). In contrast, some of the mismatching segments substantially affected the cOA production (Figure 3C). In agreement with results in Figure 2F, segment mismatches in the CAR (S1 and S2 region) completely abolished the production of cOA. Mismatches in segments S3, S5 and S6 had a minor impact on the production of cOA, whereas a major effect on cOA production was observed with mismatches in segment S4.

Since the endogenous complex is a mixture of longer and shorter complexes, we switched to using the TtCmr-46 or TtCmr-40 reconstituted complexes in order to pinpoint this crucial region more precisely (Figure 3D-I). The TtCmr-46 complex almost completely mirrored the results obtained with the endogenous complex, with the exception that mismatches in segment S4 seem to abolish the RNA targeting activity (Figure 3E). Similarly, mismatches in the CAR (S1 and S2) as well as in S4 diminish cOA production. However, this essential region appeared to have shifted one segment in the TtCmr-40 complex, with strict base pairing requirements for RNA targeting in the third and fourth segments (Figure 3H). Again, effects on cOA production mirrored these results (Figure 3I). Taken together, these results demonstrate a 3’ located seed region in TtCmr that, shifts towards the 5’ end of the crRNA with smaller crRNA sizes (in smaller TtCmr complexes). We propose that together, these regions act as flexible seed sequences in TtCmr.

### Seed region in TtCmr structure is required for concerted structural rearrangements

To determine the structural basis for the 3’ seed in RNA targeting, we interrogated our previously determined cryo-electron microscopy structures of TtCmr with both a 46 and 40-nt crRNA (EMD-2898 and −2899, respectively) [23]. In both structures, the 3’ end of the crRNA is largely exposed, as it is only cradled by the Cmr1/6 head along one side the RNA strand (Figure 4 A,B). In contrast, the seed region of the crRNA immediately upstream from the 3’ end (23 to 38 nt) is partially buried and sandwiched between the Cmr4 and Cmr5 subunits and are less exposed (Figure 4 C,D). The 3’ end of the crRNA is thus primed for transmitting conformational changes and repositioning Cmr5 subunits along the complex to facilitate complete target binding. This strongly suggests that complementarity between the crRNA and target in this region is critical for propagation of base-pairing along the length of the complex and sensitive to mismatches. Importantly, because the crRNA is shortened by one segment (6 nucleotides) in TtCmr40 compared to TtCmr46, the seed shifts towards the 5’ end of the crRNA (17 to 32 nt) in the smaller complex. Interestingly, recent structures of the *S. islandicus* Cmr complex also reveal an exposed 3’ end of the crRNA, despite the decoration of the complex by an additional 26 Cmr7 subunits that are absent in our TtCmr structure [41]. The persistent occurrence of the exposed 3’ end across Cmr complexes from multiple sources further supports our model.

**Figure 4.**
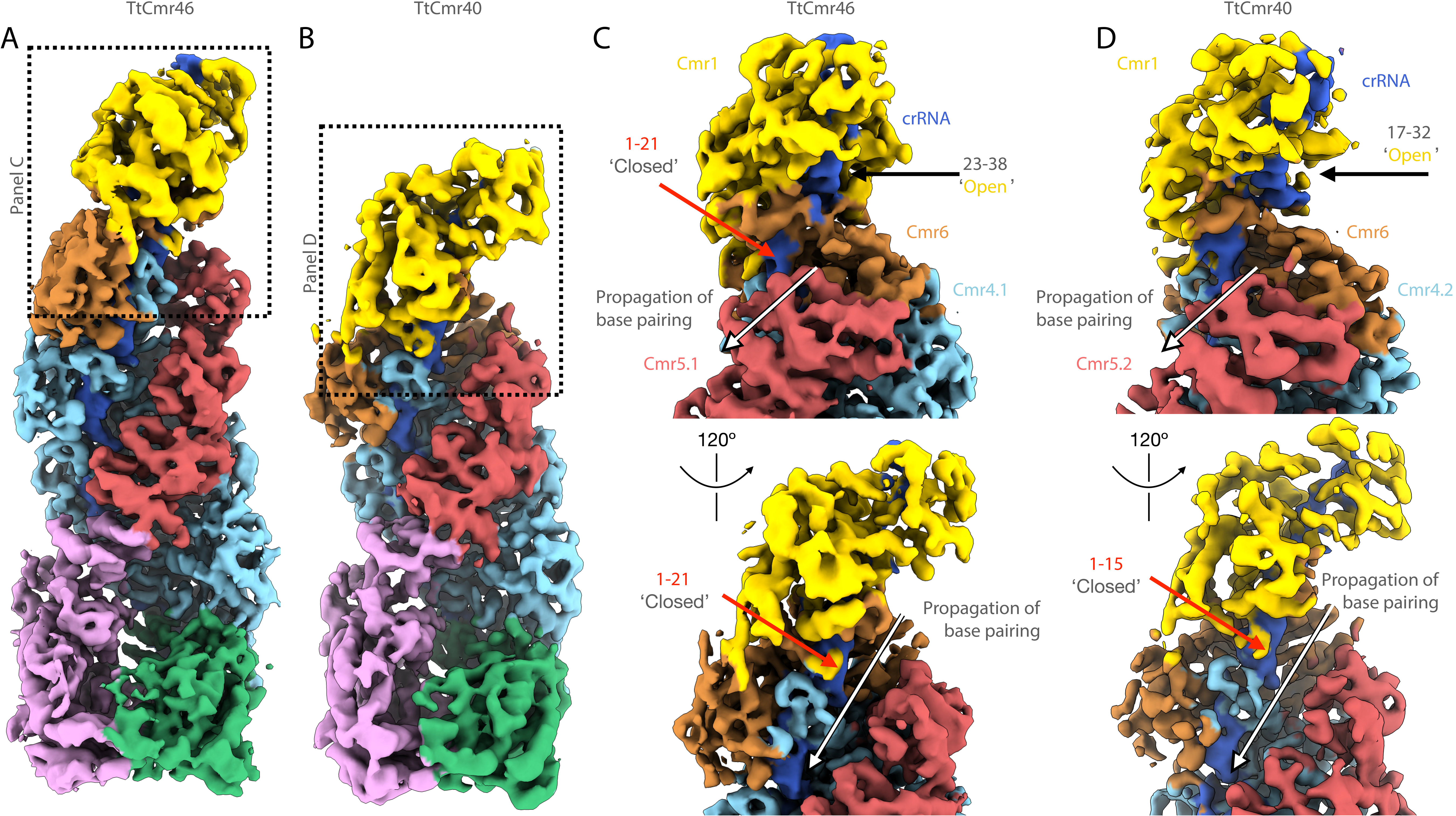
Structural basis for flexible 3’ seed region in TtCmr complexes. **(A)** Overall structure of TtCmr complex with 46 nt crRNA (EMD-2898). **(B)** Overall structure of TtCmr complex with a 40 nt crRNA (EMD-2899). Complexes are colored as in figure 1. Boxed regions refer to close-up views shown in panels C and D. **(C)** Close-up view of the top of TtCmr46. **(D)** Close-up view of the top of TtCmr34. The ‘open’ seed region located at the 3’ end of the crRNA is more accessible than the upstream regions located towards the 5’ end, which are protected by Cmr5 subunits. The 3’ end of the crRNA is thus primed for propagating conformational changes and repositioning Cmr5 subunits along the complex upon target binding.

### SCOPE - a TtCmr-based nucleic acid detection tool

Based on these stringent target RNA requirements, the need to fulfill both the seed and CAR to initiate cOA production, we hypothesized that type III CRISPR-Cas systems could be repurposed as a highly sensitive, novel nucleic acid detection tool. To investigate this possibility, we opted to couple the production of the second messenger to an easy read-out. In nature, these cOAs specifically bind to CARF domains of proteins, causing an allosteric activation of fused nuclease domains. A well-characterized example of such an enzyme is the cOA-dependent non-specific RNase (TTHB144) of *T. thermophilus* HB8 [30]. We selected this enzyme to establish a synthetic signal transduction route consisting of an RNA-targeting TtCmr/crRNA complex that generates cOA molecules, which trigger the cleavage of a reporter RNA by TTHB144 thereby generating a detectable fluorescence signal.

After visualizing the first two steps of our detection tool (target RNA binding and cOA production), we continued by monitoring the third step in the process: activation of a CARF effector protein, TTHB144. We first performed *in vitro* activity assays, using the TtCmr-46 complex, to which we added purified TTHB144 and a 5’ Cy5-labelled reporter RNA. We observed defined degradation products of the reporter RNA only when both the 4.5 target RNA (T) and TTHB144 are present (Figure 5A), whereas a non-target RNA (NT) did not induce this activity. The segment 4 (S4) mismatch target RNA (the identified seed region) greatly diminished the intensity of reporter RNA degradation products. In agreement with earlier results, target RNAs containing either mismatches in segment 1 (S1) or a single nucleotide mismatch in the CAR (C1) fully abolished TTHB144 activation, as seen by the lack of reporter RNA degradation products (Figure 5A). These results show that the target RNA sequence requirements to activate TTHB144 perfectly match with those of cOA production and that our setup can discriminate single nucleotide differences.

**Figure 5.**
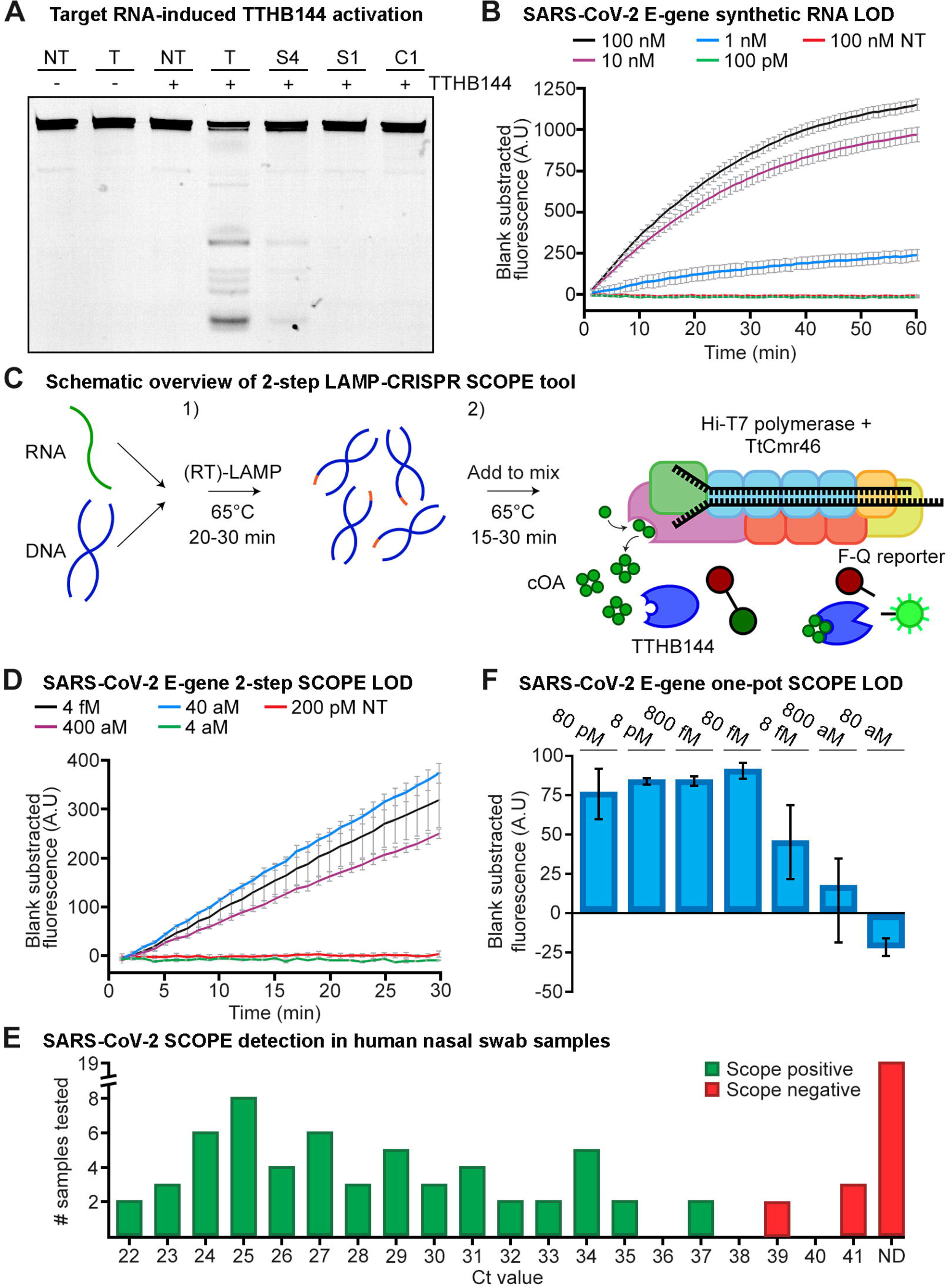
A novel type III CRISPR-Cas tool for the sensitive detection of nucleic acids. **(A)** Denaturing PAGE resulting from activity assays performed with the CARF protein TTHB144 and a 5’ Cy5 labelled reporter RNA. Activation of TTHB144 due to cOAs produced by the TtCmr complex was monitored by offering fully complementary (T) target RNAs, or target RNA with mismatches in segments one (S1), four (S4) or six (S6), or by a single mismatch in the CAR (C1). A fully non-target RNA (NT) was used as a control. **(B)** Limit of detection (LOD) assay using a synthetic SARS-CoV-2 E-gene and a fluorophore-quencher reporter RNA masking construct measured over time. A non-target RNA (NT) was used as a control. **(C)** Schematic overview of the 2-step reaction setup consisting out of (1) a (RT)-LAMP based pre-amplification step and (2) a T7-based *in vitro* transcription and type III CRISPR detection step. **(D)** Limit of detection assay using the 2-step setup (depicted in panel C), with a SARS-CoV-2 synthetic full RNA genome as target. **(E)** Detection of SARS-CoV-2 in human swab samples. Ct-values of qPCR analysis (*Orf1ab* gene) of 81 samples are depicted on the X-axis with the true negative samples displayed as not determined (ND). See Table S2 for Ct-values of qPCR analysis of SARS-CoV-2 samples and respective Scope tool score. **(F)** One-pot LAMP-CRISPR limit of detection assay on a synthetic SARS-CoV-2 E-gene.

To generate an easy read-out for our tool, we performed a similar assay, with a fluorophore-quencher reporter RNA and measured fluorescence in real-time. For this assay, we used a crRNA that targets the E-gene of the SARS-CoV-2, one of the genes used in various RT-qPCR tests for SARS-CoV-2, validated by FIND [42](Table S1). Using this setup, a sensitivity of 1 nM target RNA concentration was achieved with a fluorescence signal detectable within seconds after starting the incubation (Figure 5B).

To enable DNA detection and to enhance the sensitivity of our tool even further, we included a pre-amplification step akin to those used by other CRISPR-Cas based diagnostic tools [43]. For this proof of principle, we designed a (RT-)LAMP pre-amplification step that specifically amplifies the SARS-CoV-2 E-gene, simultaneously adding a T7 promotor to the amplicon that allowed for a subsequent *in vitro* transcription step [44] (Table S1, Figure 5C). To determine the limit of detection (LOD) of this 2-step approach, ten-fold dilutions of the SARS-CoV-2 full synthetic RNA genome (Twist Bioscience) were used. The inclusion of the LAMP pre-amplification, enhanced the sensitivity of our tool, reaching sensitivities in the attomolar (10^−18^ M) range within a timespan of ~35 minutes (30 min pre-amplification (1) + 5 min CRISPR detection (2)) (Figure 5D).

To validate our 2-step setup in a more complex reaction environment, we continued by testing human nasal swab samples, which were collected at an on-site SARS-CoV-2 testing facility. Out of the 81 samples tested, our tool scored 62 of them positively. We validated these results by comparing them to a PCR-based test (current standard for SARS-CoV-2 testing) that was performed on the same samples in parallel, which were in excellent agreement up to a relevant Ct-value of ~39. In addition, 20 samples that were considered negative by PCR (Ct value not determined, ND), were scored negative by SCOPE as well (Figure 5E).

Lastly, the thermophilic nature of the type III proteins of our tool offers an attractive opportunity for a one-pot reaction by combining the LAMP pre-amplification step with CRISPR detection, as the optimum temperature of the reactions is in the same range. Due to the maximum temperature tolerance of the Hi-T7 RNA polymerase (NEB), we performed a one-pot LAMP-CRISPR assay at a temperature of 55°C. The limit of detection was determined at 800 aM, using a synthetic version of the SARS-CoV-2 E-gene, demonstrating the feasibility of this approach (Figure 5F).

## Discussion

Recent advancements in our understanding of type III CRISPR-Cas systems have highlighted that they have unique mechanistic features compared to other CRISPR-Cas systems. Examples of this include the requirement for reverse-transcriptase activity for some type III systems during the adaptation phase [45]–[47] and the potentially large signaling network mediated by cOA molecules in the interference phase [27]–[29][32][45]–[47]. Furthermore, type III systems are exceptional in the sense that they are the only CRISPR-Cas system characterized to date capable of targeting both RNA (guide dependent) and DNA (collateral). However, the latest classification in type III CRISPR-Cas systems suggested that not all type III systems are endowed with DNase activity, due to an inactivated or missing HD domain in Cas10 [50]. This suggests that RNA is the bona fide target of these systems, as is the case for the type III-B system (Cmr-β) of S. islandicus [11], and of *T. thermophilus* presented here.

Yet another unique feature of type III systems is the variable crRNA length with a typical 6 nt periodicity, which, in turn, corresponds to the number of Cas7 subunits that constitute the backbone of type III-A and type III-B complexes [8][12][18][20][49]–[51]. Consequently, the cellular population of type III complexes are a heterogenous mixture of larger and smaller complexes [13][34]. The biological significance of these observations has remained elusive. Lastly, studies addressing the seed in type III systems are somewhat conflicting, with one report proposing a complete absence of a seed [53], whereas other studies report a seed region in either the 5’ [53][54] or 3’ end [55][56] of the crRNA guide. These discrepancies can be explained by the different methods used to pinpoint certain regions of importance on the crRNA. For instance, read-outs that either directly or indirectly look at cOA production, such as phage challenge experiments or conformational changes in Cas10, will point towards a bigger importance of the 5’ region. However, studies on the relationship of seed requirements and cOA production have paved the way for a more thorough and complete investigation [48]. Here, we dissected the importance of different size-variants of the TtCmr complex and the different regions on the crRNA by looking at each step from target RNA binding to CARF protein activation individually. Together, our data resulted in a model presented in Figure 6.

**Figure 6.**
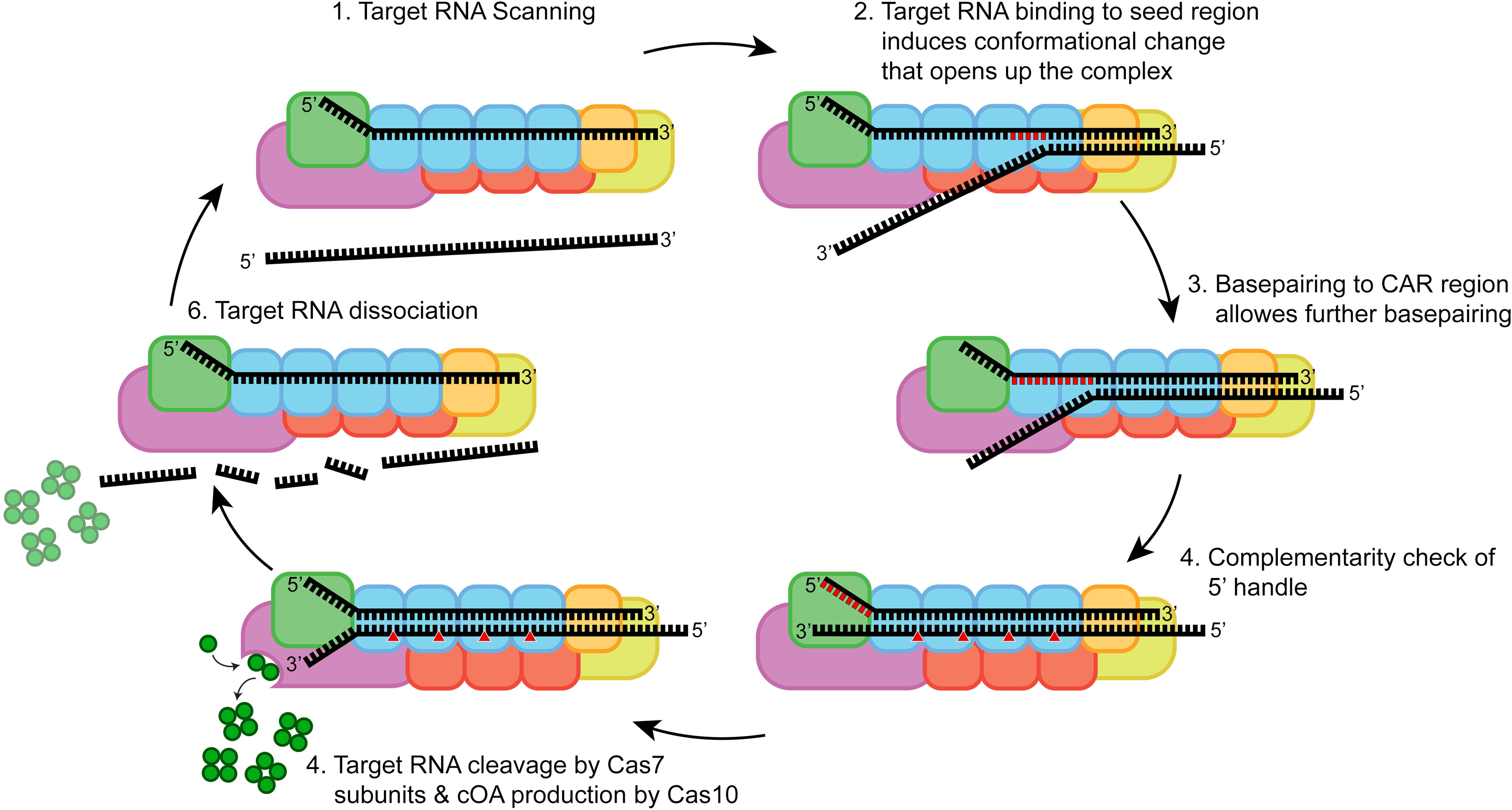
Schematic model of TtCmr target binding and subsequent activities. **(1)** TtCmr complex with bound crRNA is scanning for complimentary target RNA. **(2)** Target RNA binding is initiated at seed region, which induces a conformational change that allows further base_pairing. **(3)** Full base_pairing of target RNA to the crRNA, activating Cas10. **(4)** Target RNA is cleaved by Cas7 subunits and cOA is produced by Cas10. **(5)** Cleaved target RNA dissociates from TtCmr.

Firstly, we demonstrated that target RNA binding is initiated by a largely exposed 3’ region on the crRNA, which we designated as the seed region (Figure 3 and Figure S3). This region coincides with the first region of the crRNA leaving the Cas7/Cas11 backbone that extends to Cmr1 and Cmr6 on the top of the complex. In agreement with our observations, previous studies showed that target RNA binding in the type III-B system requires Cmr1 and Cmr6 and that base pairing in a confined region within the 3’ end of the crRNA is crucial for interference [24][55]. We therefore propose that Cmr1 and Cmr6 are involved in the proper positioning of the seed region to initiate base pairing with its cognate target RNA. The position of Cmr1 and Cmr6 on the crRNA is determined by the variable 3’ end of the crRNA. Indeed, we observed that shorter complexes (with the 40 nt crRNA) had a seed region, that was shifted one segment towards the 5’ end (Figure 3H-I). We conclude that target RNA binding (and cleavage) is governed by a flexible 3’ located seed region. This is an important difference with the structurally-related type I complexes, which harbor a fixed 5’ seed region [38].

Secondly, fulfilling the base pairing requirements in the seed region promotes a conformational change within the TtCmr complex, opening a channel along the Cas7/Cas11 backbone wide enough to accommodate further base pairing interactions of the crRNA with its target RNA (Figure 4,6). Our results show that, in contrast to target RNA degradation, cOA production is dependent on base pairing with the first seven 5’ nucleotides of the spacer part of the crRNA (with the exception of the nucleotide at position 6), a region that we designated as the CAR (Cas10 Activating Region). In accordance with previous work on type I and type III complexes, the nucleotide after each segment (i.e. every sixth nucleotide) is excluded from base pairing, due to the thumb-like extension by the Cas7 [23][58].

Lastly, even after fulfilling RNA targeting requirements (starting at the seed, followed by the CAR) we showed that a final checkpoint ensures that no cOA is produced when targeting self-RNAs (i.e. anti-sense transcripts from the CRISPR array, which has been reported to occur at low levels) [59]. Although self-RNAs are bound and cleaved by TtCmr, the complementarity between the 5’ handle and the corresponding region on the self-RNA prevented cOA production completely.

Although the biological significance of our findings still awaits further investigation *in vivo*, we anticipate that the flexibility of the seed region will complicate MGEs from escaping CRISPR-Cas targeting by introducing mutations in the protospacer. On one hand, the ability to recognize heavily mutated MGEs might be beneficial for the host to provide a robust interference response towards MGEs [59][60]. In *Marinomonas mediterranea* for example, a horizontally acquired type III-B system was recently shown to effectively bolster the immune response of its native type I-F systems to cope with phage escapees [62]. On the other hand, this same flexibility might come at the cost of a higher risk of self-targeting. While incidental binding and degradation of self-RNAs by itself might not represent a large fitness cost to the host, the subsequent activation of the Cas10 Palm domain (resulting in the production of the second messenger that allosterically activates for example promiscuous, sequence non-specific RNase activity) might have more detrimental consequences [15][16]. This scenario is in good agreement with our results, showing that activation of the Cas10 Palm domain is indeed only induced when very specific conditions are met (matching seed, matching CAR and non-matching 5’ handle).

The stringent control of cOA production prompted us to repurpose Type III CRISPR-Cas systems for the specific detection of nucleic acids. We first developed a novel pyrophosphatase-based colorimetric assay, which allows for easy quantification of oligoadenylate production. When combined with appropriate Pi calibration curves, this method could further be developed for the absolute quantification cOAs or even to quantify the virus titers in a sample. For signal amplification purposes and to achieve an easy read-out, we selected TTHB144 - one of the three native cOA-activatable (CARF) proteins present in the genome of *T. thermophilus* [27][30][63]. We observed similar trends in target RNA sequence requirements (CAR, and segments mutants) for activation of the RNase activity of TTHB144 compared to those governing cOA production (Figure 2 and 3). This shows that the stringent control of TTHB144 activation could be utilized to make distinctions between target RNAs at a single base level (Figure 5A, C1 mutant). Next to the sensitivity aspect, our tool was able to generate a signal rather quickly (detecting 1nM of target RNA within seconds, Figure 5B), due to the combination of an intrinsic signal amplification step (Cas10 producing a multitude of cOA molecules) and the high ribonuclease activity of TTHB144 [31]. For most diagnostic applications however, a sensitivity of 1nM is not high enough [43]. Therefore, we adapted a published SARS-CoV-2 LAMP pre-amplification reaction to bolster the sensitivity of our diagnostic tool even further [44]. Indeed, after testing this 2-step protocol (Figure 5C) on a SARS-CoV-2 RNA reference genome, we determined its limit of detection at 40 aM (~<1000 copies total). Validating our test on human nasal swab samples indicated our test to be 100% accurate up to a Ct value of ~39, which is higher than most accepted cutoffs for diagnosis [42]. Equally important, we did not detect any false positives out of the 20 negative samples tested.

Ideally, a true one-pot reaction is preferred to minimize steps and to reduce the risk of cross-contamination. We therefore investigated the performance of our system in a one-pot setup, at a temperature of 55°C (to comply with the upper temperature limit of Hi-T7 polymerase). Despite using non-optimal temperature for the CRISPR detection step, we were able to achieve a limit of detection of 800 aM using a synthetic gene template (Figure 5F). We showed the feasibility of the one-pot LAMP CRISPR detection approach with some room for optimization regarding the longer required incubation time. For example, the LAMP reaction by itself could be designed and optimized to work more efficiently at a temperature of 55°C by choosing a different DNA polymerase or change the primer design.

A couple of CRISPR-based nucleic acids detection platforms have been developed over the last years, such as DETECTR and SHERLOCK [64][65]. However, SCOPE is the first Class 1-based CRISPR-Cas nucleic acid detection tool, with a couple of useful characteristics (flexible seed, stringent CAR, no need for a PAM) that make it an attractive alternative over the current detection tools. Furthermore, recent bioinformatic analyses predicted that the repertoire of CARF proteins and the different catalytic activities associated with them is abundant and highly diverse, offering great opportunities for the future to utilize these activatable activities as a different readout for our tool [66].

## Acknowledgments

We thank the Nogales and Dounda labs for their support on the structural analyses (University of California, Berkeley, CA, USA). R.H.J.S was supported by a VENI grant (016.Veni.171.047), J.V.D.O. by a TOP grant (714.015.001), and S.J.J.B. by a VICI grant (VI.C.182.027), all from The Netherlands Organization for Scientific Research (NWO). This work was supported in part by Welch Foundation grant F-1938 (to D.W.T.), National Institute of General Medical Sciences (NIGMS) of the National Institutes of Health (NIH) R35GM138348 (to D.W.T.), and a Robert J. Kleberg, Jr. and Helen C. Kleberg Foundation Medical Research Award (to D.W.T.). D.W.T is a CPRIT Scholar supported by the Cancer Prevention and Research Institute of Texas (RR160088) and an Army Young Investigator supported by the Army Research Office (W911NF-19-1-0021). This work was also supported by the David Taylor Excellence Fund in Structural Biology made possible with support from Judy and Henry Sauer (to D.W.T.).

## Author Contributions

R.H.J.S., Y.Z., J.v.d.O. and J.A.S. conceived of and designed the study. Y.Z., J.A.S., D.W. T., J.P.B, S.H.P.P. and A.S. performed experiments and analyses. R.H.J.S., S.J.J.B., J.v.d.O. provided experimental guidance. R.H.J.S. and J.A.S. wrote the manuscript with significant input from other authors.

## Supplemental Information titles and legends

**Figure S1. 5’ Handle complementarity affects cOA production**

**(A)** Effects of 5’ handle complementarity on cOA production in TtCmr40 complex. **(B)** Similar to panel A, using TtCmr46 complex. NT= Non-target WT= Wild-type target

**Figure S2. Complementarity in 5’ region regulates cOA production**

**(A)** Effect of mismatches in 5’ region on production of cOA in TtCmr40 complex. **(B)** Same assay as panel A, using TtCmr46 complex instead.

**Figure S3. Base pairing of the target RNA with the TtCmr-bound crRNA is initiated at the 3’ end of the crRNA.**

**(A)** EMSA analysis of the endogenous TtCmr complex incubated with different target RNAs (table S1) each containing a stretch of 5 nt mismatching with the TtCmr-bound crRNA. **(B)** Similar EMSA analysis as panel A, using the TtCmr-46 complex. **(C)** Similar EMSA analysis as panel A, using the TtCmr-40 complex.

**Figure S4. Schematic overview of SARS-CoV-2 E-gene LAMP amplification**

Design of SARS-CoV-2 E-gene LAMP amplification, adapted from [44]. Modification to the Loop primer is the addition of a T7 RNA polymerase promoter (Table S1).

**Table S2. Ct values of qPCR analysis of SARS-CoV-2 positive samples**

## Materials and Methods

### Purification of the Cmr complex and individual subunits

A *T. thermophilus* HB8 strain was created expressing a genomic His-tagged Cm6 subunit. The (His)6-tagged TtCmr complex purification was achieved via several chromatography steps as detailed previously (Staals et al., 2013). The purified TtCmr sample was further concentrated using Vivaspin 20 centrifugal concentrator. For reconstituted complex, each subunit was individually expressed and purified. All subunits were cloned in a bicistronic design elements containing expression plasmid under an Isopropyl β-D-1-thiogalactopyranoside (IPTG) inducible T7 promoter and N-terminal Streptavidin tag. Expression plasmid containing *E. coli* BL21(DE3) were grown at 37°C until ~OD600=0.6, after which the culture was cold-shocked on ice for 1h. IPTG was added to a final concentration of 0.5-1 mM and the culture was incubated at 20°C for 16h. Cells were harvested and resuspended in Wash Buffer (150mM NaCl, 100mM Tris-HCl, pH 8) and a Complete protease inhibitor tablet was added (Roche). Cells were lysed by sonication (25% amplitude 1 sec on, 2 sec off, Bandelin Sonopuls) and spun down at 30.000g for 45 min, subsequent lysate was filter (0.45μm) clarified. A StrepTrap HP (GE) column was equilibrated using Wash Buffer and the lysate was run over it. Wash Buffer with d-Desthiobiotin added to a final concentration of 2.5 mM was used to elute. The elution fractions were pooled, concentrated and run over a HiLoad® 16/600 Superdex® 75 pg size exclusion chromatography column for further purification. Purification of TTHB144 was performed in similar fashion.

### Reconstitution of TtCmr complexes

For TtCmr46, 3.5 μL crRNA (700 ng) was added to 3.5 μL 1X Cmr buffer (20 mM Tris-HCl pH 8.0, 150 mM NaCl). Subsequently, the subunits were added to the reaction mixture in a specific order (Cmr3, Cmr2, Cmr4, Cmr5, Cmr6, Cmr1) to a final concentration of 2.5 μM, 2.5 μM, 10 μM, 7.5 μM, 2.5 μM and 2.5 μM respectively, to make up a total reaction volume of 20 μL. For TtCmr40 final concentration of subunits was adjusted to 2.5 μM, 2.5 μM, 7.5 μM, 5 μM, 2.5 μM and 2.5 μM respectively. The reaction mixture was incubated at 65°C for 30 minutes.

### In vitro cleavage activity assays

RNA substrates (listed in Table S1) were either 5’ labelled by T4 polynucleotide kinase (NEB) and 5’ 32P-γ −ATP, after which they were purified from a denaturing PAGE using RNA gel elution buffer (0.5M Sodium acetate, 10mM MgCl2, 1mM EDTA and 0.1% SDS) or ordered with a 5’ Cy5 fluorescent label. *In vitro* cleavage activity assays were conducted in TtCmr activity assay buffer (20mM Tris-HCl pH 8.0, 150mM NaCl, 10mM DTT, 1 mM ATP, and 2 mM MgCl2) using the RNA substrate and 62.5nM TtCmr. Unless stated otherwise, the reaction was incubated at 65°C for 1 hour. RNA loading dye (containing 95% formamide, dyes left out in case of Cy5 substrates) was added to the samples after incubation boiled at 95° for 5 minutes. The samples were run on a 20% denaturing polyacrylamide gel (containing 7M urea) for about 1-4 hours at 15mA or overnight at a constant of 4mA. The image was visualized using phosphorimaging or fluorescent gel scanning (IMAGER).

### cOA detection assay

The in vitro cOA detection assays were conducted in TtCmr activity assay buffer (20mM Tris-HCl pH 8.0, 150mM NaCl, 10mM DTT, 1 mM ATP, and 0.5 mM 30 MgCl2) to which Cmr-complex (62.5 nM) as well as the RNA substrates (200 nM, listed in Table S1) were added. The reaction was incubated at 65°C for 1 hour after which 0.05 units of pyrophosphatase (ThermoFisher EF0221) was added, followed by an incubation at 25°C for 30 minutes. Alternatively, thermostable pyrophosphatase (NEB #M0296) was added during the 1h incubation at 65°C. The signal was visualized using the Malachite Green Phosphate Assay Kit (Sigma-Aldrich MAK307).

### EMSA

EMSAs were performed by incubating 62.5nM TtCmr complex with 5’ Cy5 labeled target RNAs (Table S1) in Cmr binding buffer (20 mM Tris-HCl pH 8.0, 150 mM NaCl, 0.1 mM DTT, 1 mM EDTA). All reactions were incubated for 20 min at 65°C hour before electrophoresis on a native 5% (w/v) polyacrylamide gel (PAGE) at 15mA. The image was visualized via fluorescent gel scanning (GE Typhoon).

### Structural modelling

In order to model crRNA within our previously determined TtCmr46 and TtCmr40 maps (EMD-2898 and −2899, respectively), the model of *S. islandicus* Cmr complex (PD 6S6B) was fitted as a single rigid body into the TtCmr46 map, and supervised flexible fitting was performed using Isolde [67]. Maps were visualized using ChimeraX [68].

### Human swab sample collection, extraction and qPCR

PCR was conducted in a certified clinical laboratory and all procedures were validated according to the ISO 15189 standard. Nasopharyngeal swabs were transferred into 3 ml Universal transport medium. RNA was isolated and purified using the MagC extraction kit (Seegene) on an automatic nucleic acid extractor Hamilton MicroLAB StartLET (Bonaduz). SARS-CoV-2 qPCR was performed using the Allplex 19-nCoV multiplex Real-time PCR assay (Seegene), as described earlier [69]. Results were interpreted with Seegene Viewer data analysis software. A positive result was defined as amplification of the *Orf1ab* SARS-CoV-2 gene.

### Nucleic acid detection tool

The target RNA induced activation of TTHB144 assays were performed similarly to the earlier described in vitro cleavage assays. However, non-labeled target RNA was used and either a 5’ Cy5 reporter RNA or commercial RNaseAlert was added, as well as 1 μM purified TTHB144. The LAMP reaction in the 2-step LAMP-CRISPR detection was performed using the WarmStart® LAMP Kit (NEB #E1700), using primer concentration described in literature (“Rapid Detection of SARS-CoV-2 Using Reverse transcription RT-LAMP method,” 2020). A T7 promotor sequence was added to the loop primer to allow for the subsequent in vitro transcription reaction, which affected its performance only marginally when compared to the standard Loop primer (data not shown). After pre-amplification by LAMP, subsequent T7 polymerase transcription will enrich the target RNA, partly consisting of a template derived region which lies between the Loop primer and the B2 primer (Figure S4). The subsequent CRISPR detection was performed by adding 8.75 μL of CRISPR Mix (1X TtCmr Activity Assay Buffer, ~62.5 nM TtCmr complex, 500 nM TTHB144, 2 μL NTP Buffer Mix (NEB #E2050), 25 U Hi-T7 RNA polymerase (NEB #M0658), 250 nM RNaseAlert™ QC System v2 (ThermoFisher) and 8 mM final concentration MgCl). Measurements were made at 1- or 2-minute intervals at 65°C on a Bio-Rad CFX96 qPCR machine (FAM channel). The one-pot LAMP-CRISPR reaction was set up as a total reaction volume of 20 μL, according to the LAMP method described earlier with additions to the earlier described CRISPR Mix (Modification: 1 μL NTP Buffer Mix (From NEB #E2050, no additional MgCl).

