## Supplemental Information for "SCOPE: Flexible targeting and stringent CARF activation enables type III CRISPR-Cas diagnostics"

**Supplemental Information titles and legends**

**Figure S1. 5’ Handle complementarity affects cOA production**

**(A)** Effects of 5’ handle complementarity on cOA production in TtCmr40 complex. **(B)** Similar to panel A, using TtCmr46 complex. NT= Non-target WT= Wild-type target

**Figure S2. Complementarity in 5’ region regulates cOA production**

**(A)** Effect of mismatches in 5’ region on production of cOA in TtCmr40 complex. **(B)** Same assay as panel A, using TtCmr46 complex instead.

**Figure S3. Base pairing of the target RNA with the TtCmr-bound crRNA is initiated at the 3’ end of the crRNA.**

**(A)** EMSA analysis of the endogenous TtCmr complex incubated with different target RNAs (table S1) each containing a stretch of 5 nt mismatching with the TtCmr-bound crRNA. **(B)** Similar EMSA analysis as panel A, using the TtCmr-46 complex. **(C)** Similar EMSA analysis as panel A, using the TtCmr-40 complex.

**Figure S4. Schematic overview of SARS-CoV-2 E-gene LAMP amplification**

Design of SARS-CoV-2 E-gene LAMP amplification, adapted from [44]. Modification to the Loop primer is the addition of a T7 RNA polymerase promoter (Table S1).

**Table S2. Ct values of qPCR analysis of SARS-CoV-2 positive samples**

**Supplementary Figure**

**Figure S1**

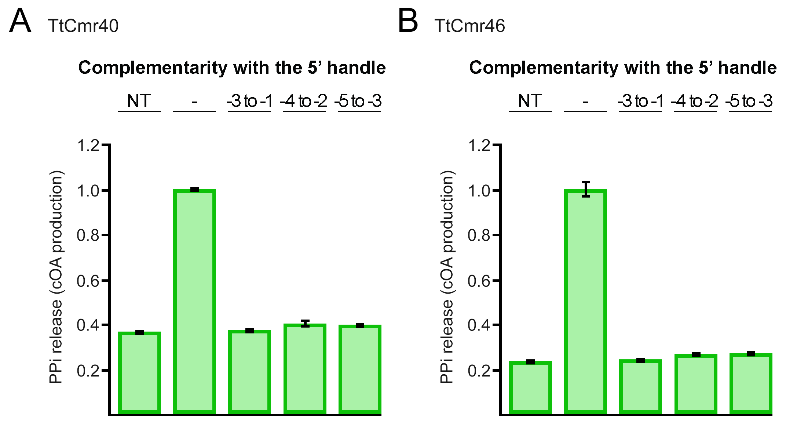

**Figure S2**

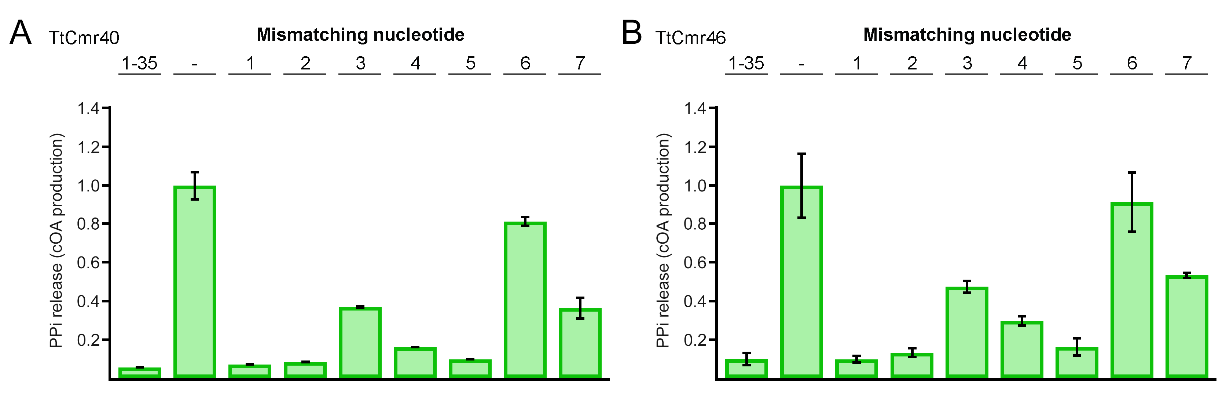

**Figure S3**

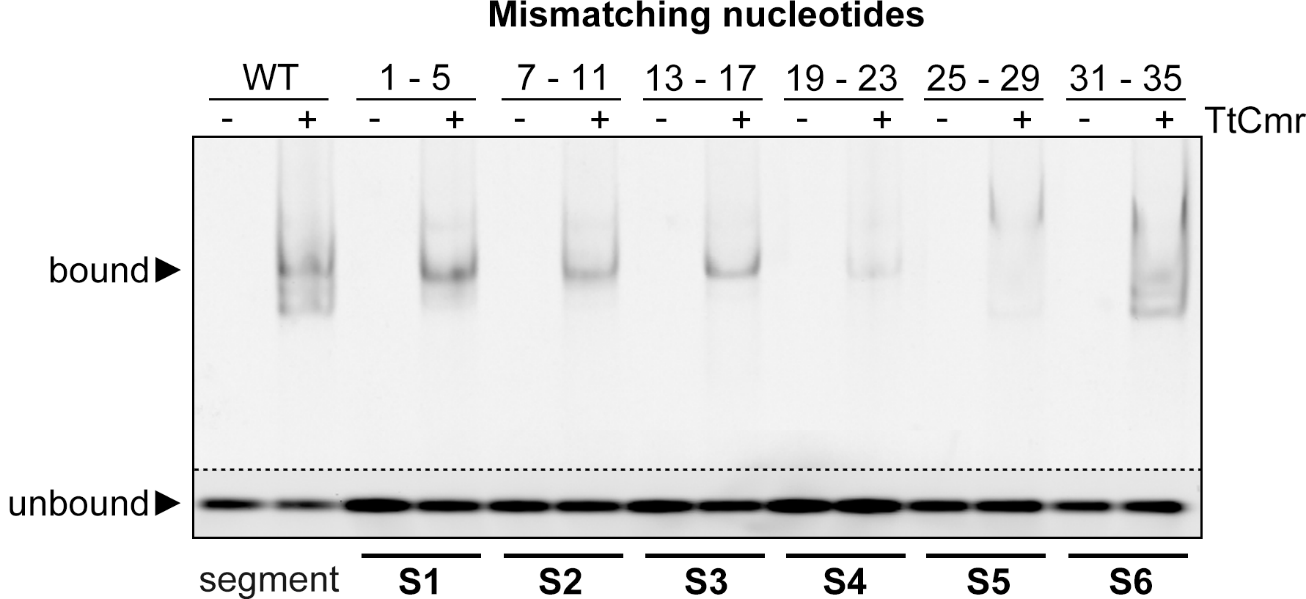

**Figure S4**

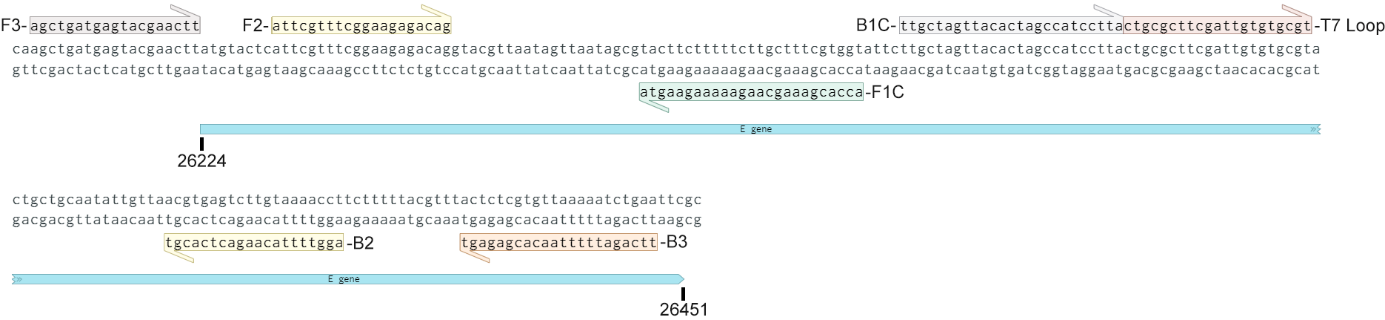

**Supplementary Table**

**Table S1**

| **Name** | **Sequence (5' to 3')** | **Comments** |
| --- | --- | --- |
| P1 | AUUGCGACCCGUAGAUAAGGCGCCCGGGGACGACCACGUCAAGGCGCAGUUGC | crRNA 4.5, 46nts (spacer region labeled as red) |
| P2 | AUUGCGACCCGUAGAUAAGGCGCCCGGGGACGAC | crRNA 4.5, 34nts (spacer region labeled as red) |
| P3 | AUUGCGACCCGUAGAUAAGGCGCCCGGGGACGACCACGUC | crRNA 4.5, 40nts (spacer region labeled as red) |
| P4 | AUUGCGACCCGUAGAUAAGGCGCCCGGGGACGACCACGUCAAGGCG | crRNA 4.5, 46nts (spacer region labeled as red) |
| P5 | GUUGUGCGCCUUGACGUGGUCGUCCCCGGGCGCCUUAUCUACGGCAGCGU | Target RNA 4.5 (red part complementary to crRNA 4.5) |
| P6 | GUUGUGCGCCUUGACGUGGUCGUCCCCGGGCGCCUUAUCUACGGGUCCGU | Matched -3/-1 target RNA (green part) |
| P7 | GUUGUGCGCCUUGACGUGGUCGUCCCCGGGCGCCUUAUCUACGGCUCGGU | Matched -4/-2 target RNA (green part) |
| P8 | GUUGUGCGCCUUGACGUGGUCGUCCCCGGGCGCCUUAUCUACGGCACGCU | Matched -5/-3 target RNA (green part) |
| P9 | GAACUGCGCCUUGACGUGGUCGUCCCCGGGCGCCUUAUCUACGCCCAUCG | Mismatched +1 RNA target (blue part) |
| P10 | GAACUGCGCCUUGACGUGGUCGUCCCCGGGCGCCUUAUCUACCGCCAUCG | Mismatched +2 RNA target (blue part) |
| P11 | GAACUGCGCCUUGACGUGGUCGUCCCCGGGCGCCUUAUCUAGGGCCAUCG | Mismatched +3 RNA target (blue part) |
| P12 | GAACUGCGCCUUGACGUGGUCGUCCCCGGGCGCCUUAUCUUCGGCCAUCG | Mismatched +4 RNA target (blue part) |
| P13 | GAACUGCGCCUUGACGUGGUCGUCCCCGGGCGCCUUAUCAACGGCCAUCG | Mismatched +5 RNA target (blue part) |
| P14 | GAACUGCGCCUUGACGUGGUCGUCCCCGGGCGCCUUAUGUACGGCCAUCG | Mismatched +6 RNA target (blue part) |
| P15 | GAACUGCGCCUUGACGUGGUCGUCCCCGGGCGCCUUAACUACGGCCAUCG | Mismatched +7 RNA target (blue part) |
| P16 | GAACUGCGCCUUGACGUGGUCGUCCCCGGGCGCCUUAUCAUGCCCCAUCG | Mismatched +1-5 RNA target (blue part), also used as 5' Cy5 variant |
| P17 | GAACUGCGCCUUGACGUGGUCGUCCCCGGGCGCGAAUACUACGGCCAUCG | Mismatched +7-11 RNA target (blue part), also used as 5' Cy5 variant |
| P18 | GAACUGCGCCUUGACGUGGUCGUCCCCCCCGCCCUUAUCUACGGCCAUCG | Mismatched +13-17 RNA target (blue part), also used as 5' Cy5 variant |
| P19 | GAACUGCGCCUUGACGUGGUCCAGGGCGGGCGCCUUAUCUACGGCCAUCG | Mismatched +19-23 RNA target (blue part), also used as 5' Cy5 variant |
| P20 | GAACUGCGCCUUGACCACCACGUCCCCGGGCGCCUUAUCUACGGCCAUCG | Mismatched +25-29 RNA target (blue part), also used as 5' Cy5 variant |
| P21 | GAACUGCGCGAACUCGUGGUCGUCCCCGGGCGCCUUAUCUACGGCCAUCG | Mismatched +31-35 RNA target (blue part), also used as 5' Cy5 variant |
| P22 | AAACGACGGCCAGUGCCAAGCUUACUAUACAACCUACUACCUCAU | TTHB144 5' Cy5 reporter RNA |
| P23 | AUUGCGACCACACAAUCGAAGCGCAGUAAGGAUGGCUAGUGUAACU | crRNA SARS-CoV-2 E-gene, 46nts (spacer region labelled as red) |
| P24 | UAGUUACACUAGCCAUCCUUACUGCGCUUCGAUUGUGUGCGUACUGCUGCAAUAUUGUUA | Target RNA SARS-CoV-2 E-gene (red part complementary to P23) |
| P25 | AUUGCGACACAAUAUUGCAGCAGUACGCACACAAUCGAAGCGCAGA | crRNA SARS-CoV-2 E-gene LAMP (spacer region labelled as red) |
| P26 | AGCTGATGAGTACGAACTT | LAMP E-gene F3 |
| P27 | TTCAGATTTTTAACACGAGAGT | LAMP E-gene B3 |
| P28 | ACCACGAAAGCAAGAAAAAGAAGTATTCGTTTCGGAAGAGACAG | LAMP E-gene FIP |
| P29 | TTGCTAGTTACACTAGCCATCCTTAGGTTTTACAAGACTCACGT | LAMP E-gene BIP |
| P30 | TAATACGACTCACTATAGCTGCGCTTCGATTGTGTGCGT | LAMP E-gene T7 Loop |

**Table S2**

| **Sample Name** | **Ct qPCR Orf1ab** | **qPCR result** | **Δsignal Scope** | **Scope result** |
| --- | --- | --- | --- | --- |
| Negative control | ND | Neg | 164 | Neg |
| UMCU sample 01 | 29.6 | Pos | 5664 | Pos |
| UMCU sample 02 | 36.0 | Pos | 6199 | Pos |
| UMCU sample 03 | 39.4 | Pos | 156 | Neg |
| UMCU sample 04 | 26.2 | Pos | 6526 | Pos |
| UMCU sample 05 | 32.6 | Pos | 6019 | Pos |
| UMCU sample 06 | 24.9 | Pos | 6756 | Pos |
| UMCU sample 07 | ND | Neg | 161 | Neg |
| UMCU sample 08 | 41.6 | Pos | 170 | Neg |
| UMCU sample 09 | 26.6 | Pos | 6828 | Pos |
| UMCU sample 10 | 27.4 | Pos | 6417 | Pos |
| UMCU sample 11 | ND | Neg | 155 | Neg |
| UMCU sample 12 | 25.0 | Pos | 6244 | Pos |
| UMCU sample 13 | ND | Neg | 152 | Neg |
| UMCU sample 14 | ND | Neg | 160 | Neg |
| UMCU sample 15 | 41.8 | Pos | 176 | Neg |
| UMCU sample 16 | 25.9 | Pos | 6371 | Pos |
| UMCU sample 17 | 30.9 | Pos | 6290 | Pos |
| UMCU sample 18 | 26.0 | Pos | 6394 | Pos |
| UMCU sample 19 | 31.6 | Pos | 6372 | Pos |
| UMCU sample 20 | 25.0 | Pos | 6563 | Pos |
| UMCU sample 21 | 25.6 | Pos | 6057 | Pos |
| UMCU sample 22 | 28.4 | Pos | 6058 | Pos |
| UMCU sample 23 | ND | Neg | 370 | Neg |
| UMCU sample 24 | 29.2 | Pos | 6095 | Pos |
| UMCU sample 25 | 27.7 | Pos | 6032 | Pos |
| UMCU sample 26 | 26.6 | Pos | 5979 | Pos |
| UMCU sample 27 | ND | Neg | 190 | Neg |
| UMCU sample 28 | 34.2 | Pos | 6671 | Pos |
| UMCU sample 29 | 34.7 | Pos | 6390 | Pos |
| UMCU sample 30 | ND | Neg | 174 | Neg |
| UMCU sample 31 | 28.6 | Pos | 6468 | Pos |
| UMCU sample 32 | 29.3 | Pos | 6312 | Pos |
| UMCU sample 33 | 31.3 | Pos | 6367 | Pos |
| UMCU sample 34 | 28.5 | Pos | 7056 | Pos |
| UMCU sample 35 | 30.2 | Pos | 7488 | Pos |
| UMCU sample 36 | ND | Neg | 160 | Neg |
| UMCU sample 37 | ND | Neg | 175 | Neg |
| UMCU sample 38 | ND | Neg | 184 | Neg |
| UMCU sample 39 | 25.3 | Pos | 6493 | Pos |
| UMCU sample 40 | 24.7 | Pos | 6388 | Pos |
| UMCU sample 41 | 34.9 | Pos | 5912 | Pos |
| UMCU sample 42 | ND | Neg | 200 | Neg |
| UMCU sample 43 | 30.9 | Pos | 6386 | Pos |
| UMCU sample 44 | 27.3 | Pos | 6938 | Pos |
| UMCU sample 45 | 26.8 | Pos | 6940 | Pos |
| UMCU sample 46 | 32.3 | Pos | 6809 | Pos |
| UMCU sample 47 | 35.5 | Pos | 7240 | Pos |
| UMCU sample 48 | 24.7 | Pos | 5565 | Pos |
| UMCU sample 49 | ND | Neg | 167 | Neg |
| UMCU sample 50 | 41.8 | Pos | 195 | Neg |
| UMCU sample 51 | 29.1 | Pos | 6360 | Pos |
| UMCU sample 52 | 23.0 | Pos | 6146 | Pos |
| UMCU sample 53 | 25.5 | Pos | 5731 | Pos |
| UMCU sample 54 | ND | Neg | 221 | Neg |
| UMCU sample 55 | 37.0 | Pos | 5973 | Pos |
| UMCU sample 56 | 25.5 | Pos | 6972 | Pos |
| UMCU sample 57 | 31.8 | Pos | 6698 | Pos |
| UMCU sample 58 | 37.6 | Pos | 7093 | Pos |
| UMCU sample 59 | 24.1 | Pos | 7426 | Pos |
| UMCU sample 60 | 39.4 | Pos | 150 | Neg |
| UMCU sample 61 | 24.5 | Pos | 6269 | Pos |
| UMCU sample 62 | 33.9 | Pos | 6213 | Pos |
| UMCU sample 63 | ND | Neg | 165 | Neg |
| UMCU sample 64 | 27.911 | Pos | 6262 | Pos |
| UMCU sample 65 | ND | Neg | 221 | Neg |
| UMCU sample 66 | ND | Neg | 185 | Neg |
| UMCU sample 67 | 23.6 | Pos | 6865 | Pos |
| UMCU sample 68 | 22.9 | Pos | 6938 | Pos |
| UMCU sample 69 | 27.2 | Pos | 7046 | Pos |
| UMCU sample 70 | 29.6 | Pos | 7241 | Pos |
| UMCU sample 71 | 23.9 | Pos | 7802 | Pos |
| UMCU sample 72 | 27.5 | Pos | 5164 | Pos |
| UMCU sample 73 | ND | Neg | 157 | Neg |
| UMCU sample 74 | 23.8 | Pos | 6287 | Pos |
| UMCU sample 75 | 34.2 | Pos | 6314 | Pos |
| UMCU sample 76 | 31.3 | Pos | 6438 | Pos |
| UMCU sample 77 | 34.1 | Pos | 5700 | Pos |
| UMCU sample 78 | 24.9 | Pos | 6865 | Pos |
| UMCU sample 79 | 33.4 | Pos | 6832 | Pos |
| UMCU sample 80 | ND | Neg | 200 | Neg |
| Positive control | 31.1 | Pos | 7205 | Pos |
| Positive control 2 | 35.5 | Pos | 7172 | Pos |
